# Leveraging mutational burden for complex trait prediction in sorghum

**DOI:** 10.1101/357418

**Authors:** Ravi Valluru, Elodie E. Gazave, Samuel B. Fernandes, John N. Ferguson, Roberto Lozano, Pradeep Hirannaiah, Tao Zuo, Patrick J. Brown, Andrew D.B. Leakey, Michael A. Gore, Edward S. Buckler, Nonoy Bandillo

## Abstract

Sorghum (*Sorghum bicolor* (L.) Moench) is a major staple food cereal for millions of people worldwide. The sorghum genome, like other species, accumulates deleterious mutations, likely impacting its fitness. Though selection keeps deleterious mutations rare, their complete removal from the genome is impeded due to lack of recombination, drift, and their coupling with favorable loci. To study how deleterious mutations impact agronomic phenotypes, we identified putative deleterious mutations among ~5.5M segregating variants of 229 diverse sorghum lines. We provide the whole-genome estimate of the deleterious burden in sorghum, showing that about 33% of nonsynonymous substitutions are putatively deleterious. The pattern of mutation burden varies appreciably among racial groups; the *caudatum* shows higher mutation burden while the *guinea* has lower burden. Across racial groups, the mutation burden correlated negatively with biomass, plant height, Specific Leaf Area (SLA), and tissue starch content, suggesting deleterious burden decreases trait fitness. Putatively deleterious variants explain roughly half of the genetic variance. However, there is only moderate improvement in total heritable variance explained for biomass (7.6%) and plant height (5.2%). There is no advantage in total heritable variance for SLA and starch. The contribution of putatively deleterious variants to phenotypic diversity therefore appears to be dependent on the genetic architecture of traits. Overall, our results suggest that including putatively deleterious variants in models do not significantly improve breeding accuracy because of extensive linkage. However, knowledge of deleterious variants could be leveraged for sorghum breeding through genome editing.

## INTRODUCTION

Plant genomes continually accumulate new mutations due to population demographic history [1], random drift [2], the mating system [3], domestication [4,5], and linked selection due to genetic interactions [6,7]. While a sizeable portion of such new mutations are neutral [8,9], a small portion of mutations are likely to be deleterious because they disrupt evolutionarily conserved sites, protein function [10,11], or gene expression [12] in a way that results in negative impacts on fitness. Therefore, the elimination of deleterious mutations from breeding populations has been suggested as a prospective avenue for crop improvement [13].

Sorghum ( *Sorghum bicolor* (L.) Moench, 2n = 20) is an important and versatile crop that is grown for food, forage, and fuel. It was domesticated from its wild ancestor about 8,000 years ago in Africa [14]. Five major morphological forms have traditionally been recognized, namely *bicolor, caudatum, durra, guinea,* and *kafir*. While these races are widespread in distinct regions of Africa reflecting the diverse agro-eco-environments [15,16], sorghum has maintained minimal genome redundancy due to the absence of any whole genome duplication for over 70 million years [17,18]. However, inbreeding sorghum is likely to accumulate more slightly deleterious mutations when compared to an outcrossing species, which accumulates strong recessive deleterious mutations that reduce the mean fitness of the population over time [13]. Nonetheless, there is accumulating evidence for the impact of enhanced homozygosity [19], relaxed selection [20], and low levels of outcrossing [21,22] on the frequency of deleterious polymorphisms in selfing populations. Although the relative contributions of these processes to mutation load has long been debated, both theoretical and experimental evidence suggests that reduced population size effects usually outcompete processes that enhance purging of deleterious mutations caused by selfing [20,23–25] leading to an influx of deleterious mutations into selfing species.

Modern breeding and domestication results in an increased genetic load in domesticates when compared to their wild progenitors, and a decreased load in elite cultivars when compared to landraces [4,26]. The demographic history and inbreeding allow deleterious variants of weaker effect to reach appreciable frequencies owing to random drift, which can contribute significantly to mutation load and affect fitness-related traits [27]. An estimated 20 to 30% of nonsynonymous variants are deleterious in rice [5], Arabidopsis [28], maize [29], and cassava [4]. Renaut and Rieseberg [30] identified an excess of nonsynonymous Single Nucleotide Polymorphisms (SNPs) segregating in domesticated sunflower and globe artichoke relative to natural populations. Similarly, 20 to 40% of protein-coding SNPs are predicted to have a deleterious allele in maize [29]. Indeed, deleterious mutations are predicted to be enriched near regions of strong selection [26,27], pointing to a potentially important role for deleterious variants in shaping agronomic phenotypes.

Genomic Selection (GS) can accelerate crop breeding when compared to conventional phenotypic selection approaches. In the Genome-Wide Prediction (GWP) models employed in GS, the genetic variance is modeled by accounting for either the biological additive and dominant effects of the markers, which improves the accuracy of predictions [31,32]. Genes associated with complex traits carry an uncertain number of deleterious mutations distributed across the genome, and such a mutational load may significantly contribute to the total phenotypic variation of traits [33]. Because deleterious mutations can occur in both homozygous and heterozygous states depending on the genetic context, trait-specific and genetic-context based GWP models can capture the effects of deleterious mutations. Therefore, GWP models encompassing deleterious mutations are expected to account for the total genetic contribution to, and improve the prediction ability of, complex traits [33]. However, the improvement of GWP will depend on how strongly correlated deleterious variants are to all other variants.

In this study, we examine the contribution of putatively deleterious variants to phenotypic variation in sorghum. We used a racially, geographically, and phenotypically diverse biomass sorghum population that represents the ancestry of five major sorghum types [34]. All accessions were phenotyped for two agronomic traits, dry biomass (DBM) and plant height (PHT), and for two physiological traits, specific leaf area (SLA) and tissue starch content (TSC) under field conditions. We performed whole-genome resequencing (WGS) on 229 sorghum lines and identified genome-wide putative deleterious mutations. Our main objectives of this study were to determine (1) whether empirical patterns of deleterious mutational burden differ among sorghum racial groups; and (2) whether deleterious variants improve prediction ability of complex traits, and if so, whether such abilities differ over phenotypic traits that have different genetic architecture. To address these questions, we first identified the genome-wide putative deleterious mutations and their biological effect sizes and then, estimated an individual mutation burden and its effect on phenotypic traits. Next, taking advantage of a Bayesian genomic selection framework [35], we tested the biological significance of deleterious variants in the prediction of DBM, PHT, SLA, and TSC.

## RESULTS

### Identification of putatively deleterious mutations

We resequenced the whole genome of 229 diverse biomass sorghum to an average depth of 4X and identified ~5.5M segregating variants (see Methods), of which 6.3% are located in coding regions. To determine the distribution of deleterious mutations in the sorghum genome, we first annotated deleterious variants using a SIFT score (SIFT<0.05) that predicts an amino acid substitution effect on protein function [36]. Approximately 33% of the total nonsynonymous substitutions are putatively deleterious (average SIFT score of 0.08), while 67% are predicted as tolerant mutations (average SIFT score of 0.47). The majority (75%) of nonsynonymous deleterious mutations had an average SIFT score of <0.01 (Fig. S1a). All identified deleterious mutations show comparably similar distributions among all chromosomes ( *P* = 0.34; Fig. S1b) and arise from noncentromeric regions of the chromosomes (Fig. S2). Consistent with population genetic expectations, all deleterious mutations show a low overall allele frequency (average MAF=0.07, Fig. S1c).

We then estimated the derived allele frequency (DAF) spectrum based on ‘derived deleterious allele’ which is defined as a minor allele among multi-species alignment [33]. This revealed that a large proportion of deleterious mutations have a lower DAF (<0.05)(Fig 1a). While DAF is strongly negatively associated with GERP scores [33](Fig 1b), it is positively associated with SIFT scores (Fig. S3). These results corroborate previous studies showing that selection acts to keep deleterious mutations rare [29], and support the proposition for a combined use of SIFT and GERP scores as quantitative measures of an observed variant for its long-term fitness consequences [33].

**Figure 1.**
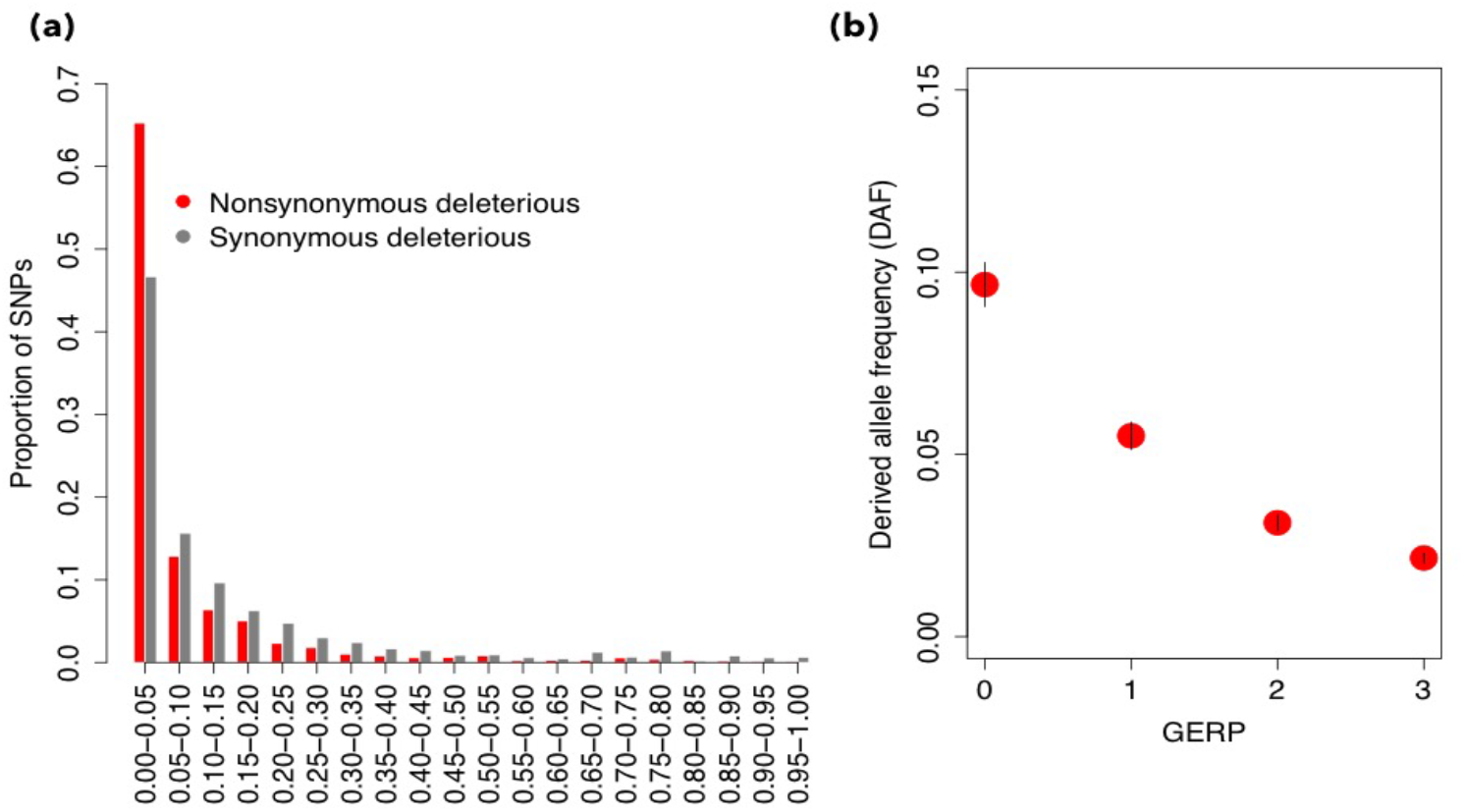
Deleterious mutations in the sorghum genome. **(a)** Site allele-frequency spectrum of deleterious mutations in the sorghum genome. The Derived Allele Frequency (DAF) distribution of alleles is shown where a minor allele from multi-species alignment was considered as a derived allele [33]. **(b)** The allele frequency of the derived alleles in bins of different GERP score.

### Distribution of effect sizes for deleterious variants

For each phenotype, we estimated the additive effect sizes explained by both deleterious variants (High-GERP deleterious variants; hereafter called HGERP_DEL-SNPs_) and random variants that are not in linkage disequilibrium (LD) but have similar allele frequency range of deleterious variants across the genome. We compared the full density distribution of the effect sizes of both deleterious and random variants to avoid the winner’s curse [37,38], and examined whether deleterious variants effect sizes are overall larger in magnitude than random variants (Fig 2).

**Figure 2.**
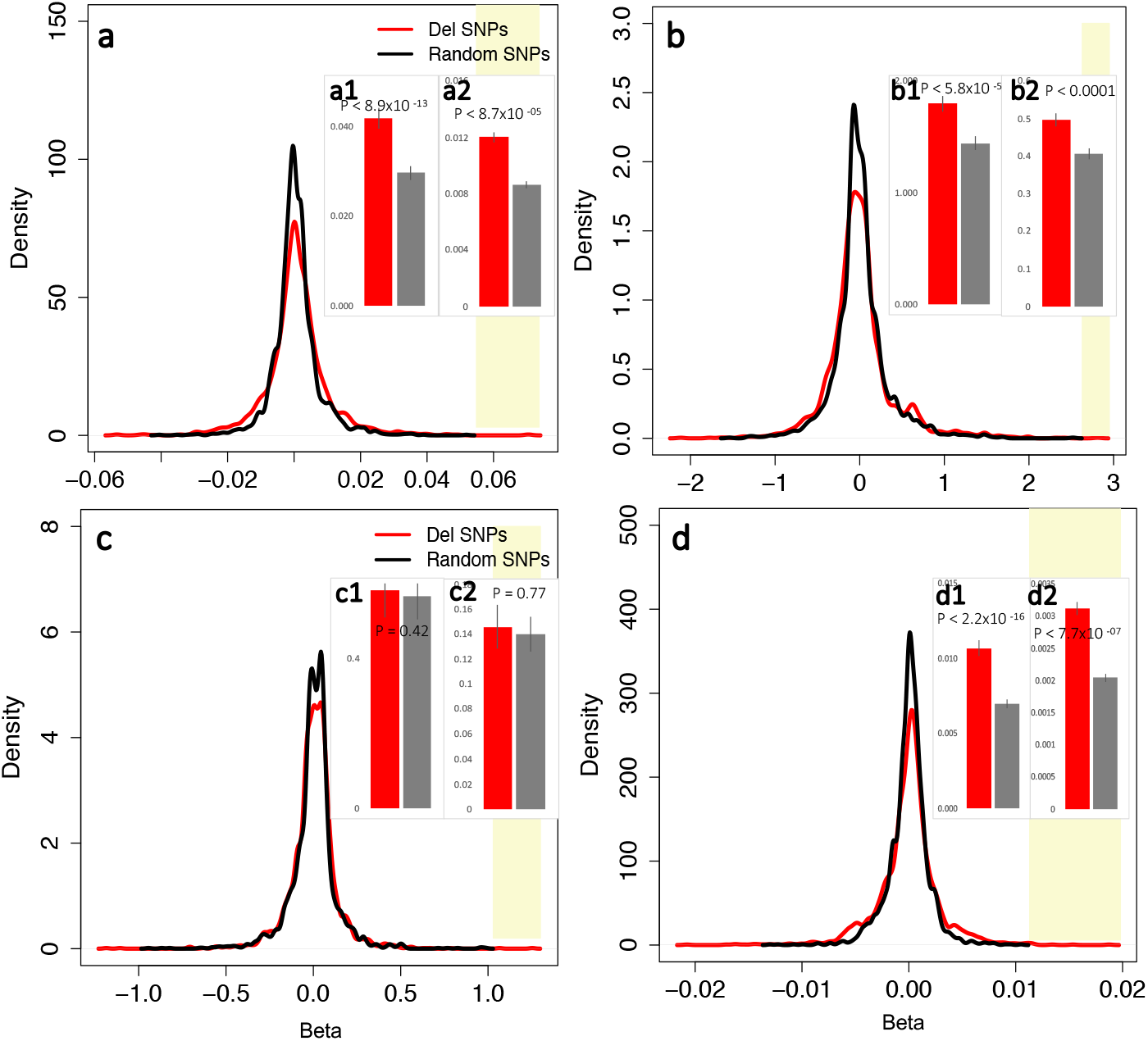
Smoothed estimate of density distribution of regression coefficients associated with deleterious variants and common variants for phenotypic traits ((a) biomass, (b) plant height, (c) Specific Leaf Area (SLA), (d) tissue starch content) for deleterious variants (red) and random variants (black) for high GERP deleterious variants (HGERP_DEL-SNPs_). The yellow vertical band indicate the extent of an additional density distribution of deleterious variants as compared to the density distribution of random variants. Inset: Barplot of the effect sizes of deleterious variants and nondeleterious variants based on the top 1% and 25% for biomass (a1,a2), plant height (b1,b2), SLA (c1,c2), and starch (d1,d2).

Our results indicate that the density distribution of the effect sizes of both deleterious and random variants show slightly different patterns. The density distribution of deleterious variants extends much farther than the distribution of random variants at the highest range (Fig 2)[37]. Such a density distribution could be due to a reduced density peak for deleterious variants; however, it was consistently observed for all traits (Fig. 2a-d). When compared at the same threshold (top 1% and 25%), deleterious variants have significantly higher average effect sizes of 30.4% (Fig. 2a1-d1) and 26.7% (Fig. 2a2-d2) across four traits over nondeleterious variants, respectively (Fig 2, Inset). Plant height has the largest effect deleterious variants as compared to other traits. These results suggest that a minor proportion of deleterious variants appear to have larger biological effect sizes when compared to nondeleterious variants that could cumulatively affect phenotypes. This is consistent with recent studies showing rare deleterious variants having much greater effect sizes than those of common variants in maize [33], human [38,39], and mouse [40].

### Deleterious mutation burden and its effect on phenotypes

We estimated the burden of deleterious variants as the count of deleterious variants corrected for the number of variants scored in all genotypes of the population (Fig. 3). Our burden estimation reveals a substantial variation for mutation burden among racial groups ( *P* = 3.14 × 10-05) under the HGERPDEL-SNPs model (Fig. 3). We observed that *caudatum* is significantly enriched, with an average of 36%, for homozygous mutation burden as compared to other racial groups. Compared to the median burden across all racial groups, *guinea* has a proportionately lower burden (−20%), while *caudatum* has a proportionately higher burden (+49%). On average, an individual typically carries 0.0112 (s.d. 0.006), 0.0124 (s.d. 0.006), 0.0140 (s.d. 0.006), and 0.0178 (s.d. 0.007) mutation burden in the homozygous state in the *guinea*, *durra*, *kafir* and *caudatum* groups, respectively. Across all racial groups, individual mutation burden ranges from 0.001 to 0.038 under the HGERP_DEL-SNPs_ model,suggesting that all racial groups showed variable mutation burden.

**Figure 3.**
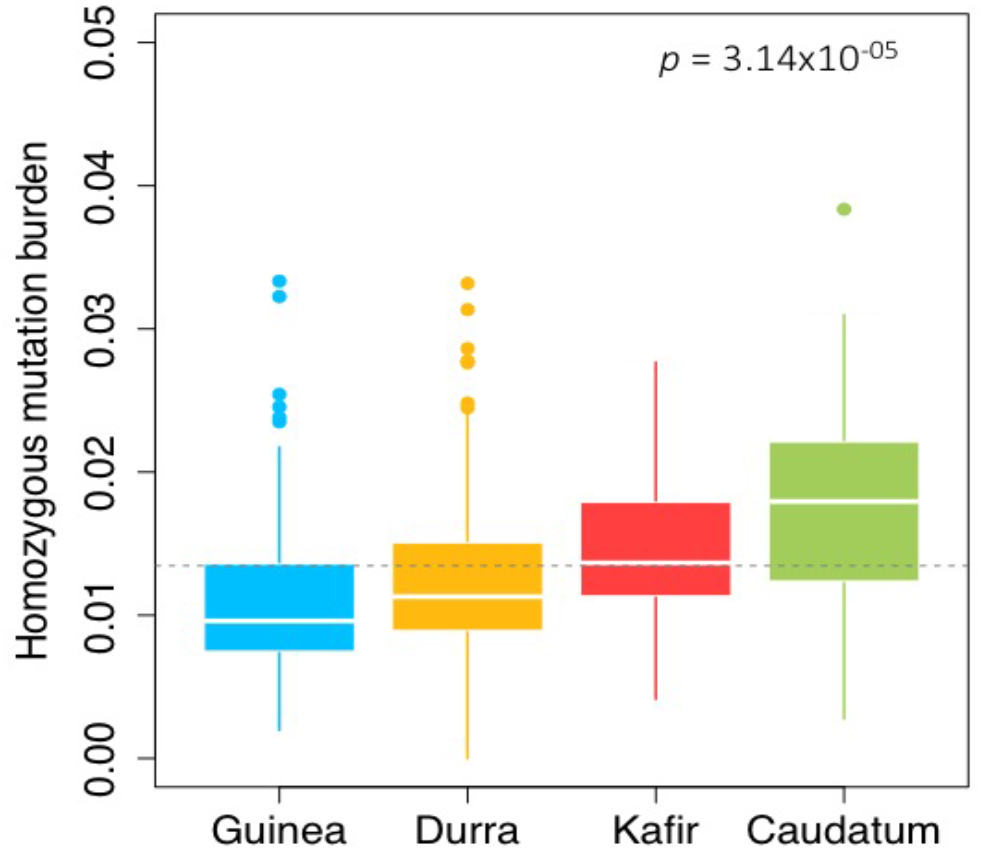
Homozygous mutation burden estimated for different racial groups of sorghum under an evolutionary-based genomic model (HGERP_DEL-SNPs_). The derived allele is defined as a minor allele from multi-species sequence alignments [33]. The total number of homozygous deleterious alleles identified within each individual was corrected for the total number of variants scored within each individual in order to avoid any bias in burden among individuals due to missing genotypic data. The horizontal broken line indicates the mean of burden across all groups.

We further evaluated the underlying relationship of mutation burden with phenotypic traits. Four phenotypic traits were selected for this study: dry biomass, plant height, SLA, and tissue starch content. We observed a substantial phenotypic variation for all traits among racial groups (Fig. S4, biomass: *P* <0.001; height: *P* <0.05; SLA: *P* <0.001; starch: *P* <0.05), with highly heritable variation observed for plant height (*H^2^*=0.87) and biomass (*H^2^*=0.73), consistent with previous studies [34]. We also found strong racial group-specific correlations among traits (Fig. S5). Using a simple linear regression model between mutation burden and phenotypic traits, across all racial groups, we consistently found a negative relationship of mutation burden with all traits (Table S1), suggesting that deleterious variants decrease trait fitness.

### Accounting for deleterious variants in genome-wide prediction

We tested whether incorporating putatively deleterious variants could inform genomic selection (GS) models and improve phenotype prediction. Deleterious mutations identified from WGS were used as priors and integrated into a genomic prediction framework (Fig. 4). We quantified the amount of genetic variance, heritability, and model improvement by deleterious variants and compared with that of random variants. Based on a variance partitioning approach with a two-kernel model (see Methods), the model with putatively deleterious variants explained roughly half of the genetic variance (biomass: 52%, plant height: 54%, SLA: 48%, and starch: 46%) (Fig. 4a). However, there was only a moderate improvement in total heritable variance explained for biomass (7.6%, h2 = 0.24 against 0.22 for random variants) and plant height (5.2%, h2 = 0.36 against 0.34 for random variants), and no advantage for SLA and starch (Fig. 4b) as compared to random variants.

**Figure 4.**
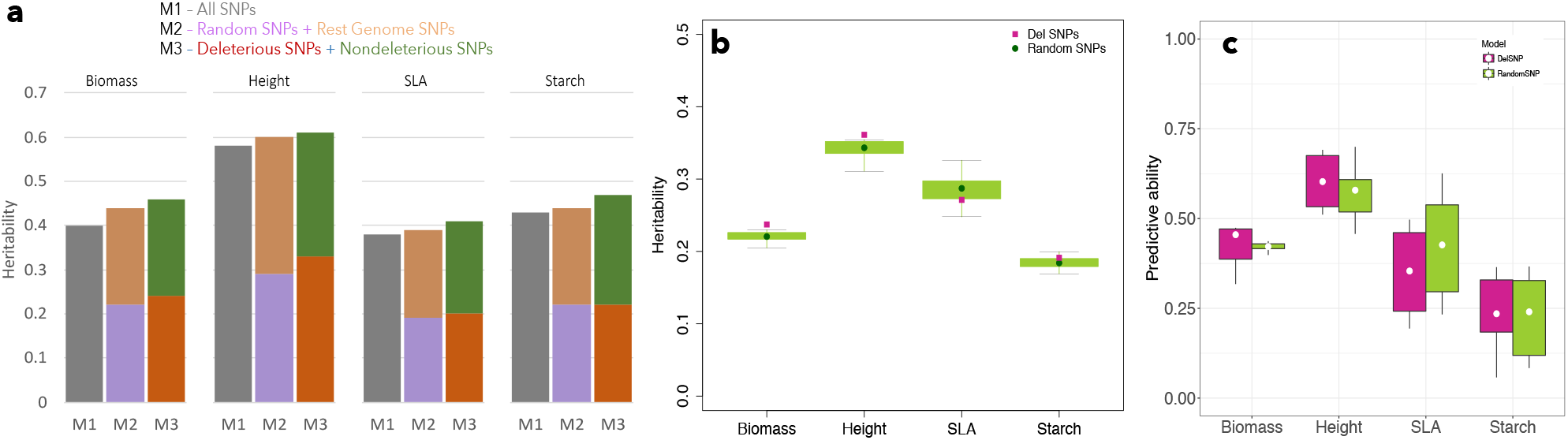
Genome-wide prediction models incorporating putatively deleterious mutations. (a-b) Heritability estimates for all four traits using either a (a) two-kernel model or (b) single-kernel model. Heritability estimates for random variants are derived based on 100 independent sets that are randomly chosen across the genome from variants that are not in LD with deleterious variants. (c) Boxplots showing a five-fold cross validation prediction ability estimation for deleterious variants and random variants.

To evaluate the predictive ability, we performed a five-fold cross-validation implemented in a GBLUP model with either the deleterious or the random SNP data sets. Consistent with the results of heritability, we observed an 8.1% and 4.2% improvement on predictive ability for biomass and plant height, respectively, while there was no improvement for SLA and starch (Fig. 4c). These results suggest that the contribution of putatively deleterious variants to phenotypic variation varies considerably among traits.

## DISCUSSION

Sorghum, a genus that evolved across diverse environments in Africa, exhibits a wide range of phenotypic diversity [15,41,42]. This raises the question of whether sorghum racial groups carry variable deleterious load, allowing the mutation consequences to be tested for phenotypic diversity. In this study, we WGS 229 sorghum lines and defined genome-wide putative deleterious mutations using SIFT and GERP scores. All racial groups of sorghum showed variable mutation burden (ranged from 0.001-0.038) that correlated negatively with phenotypic traits. We observed that a minor proportion of deleterious mutations had larger biological effects. We further noticed that the prediction ability of the genome-wide prediction models encompassing deleterious variants are largely trait-dependent.

Combining the criteria of SIFT and GERP scores, we first show that sorghum racial groups accumulate considerable amounts of deleterious mutations in the genome, estimated to be ~33% of total nonsynonymous substitutions (Fig. 1). Although the number and frequency of such mutations within a population depends on population size, our results match well with previous studies that estimated 20 to 30% of nonsynonymous variants to be deleterious in several crop species, including model plant species [4,5,28,29]. Considering only highly conserved (GERP>2) and frequent (DAF>0.9) mutations, there are 63 nonsynonymous deleterious mutations across racial groups, and distributed across all chromosomes. These variants are likely a combination of variants of important domestication targets, recent pseudogenes, and some truly deleterious variants that are the product of drift.

We next estimated an individual mutation burden as the count of deleterious variants corrected for the number of variants scored, which differed considerably among individuals and racial groups (Fig. 3). It is notable but expected given that different racial groups have had varying patterns of population dynamics, selection intensities, and domestication histories that could alter the influx of deleterious mutations [14,15,17]. Contrasting deleterious burden has previously been reported in different populations of crop species [4,5,30], and humans [43–45]. Comparatively, the *caudatum* group appears to have a higher mutation burden than the *guinea* group; the oldest of the specialized sorghum races [46,47]. We propose that the higher mutation burden of the *caudatum* group might be potentially related to the population bottleneck, resulting in a smaller population size that increases the chances of inbreeding, genetic homogeneity, and an increased influx of deleterious mutations [13,30,33]. On the other hand, a lower mutation burden in the *guinea* might be due to its higher outcrossing rates, which can reach up to 20% when compared to other races [48,49]. Our results, therefore, suggest that, first, negative selection is less effective at removing weakly deleterious mutations, yielding variable mutation burden among racial groups. Second, the combined effects of a bottleneck and directional selection during domestication [43,50] can have an important impact on the deleterious burden even in smaller racial groups of sorghum in which founder events can be more frequent [51,52].

Although informative, our estimation of mutation burden has some important limitations. First, the deleterious mutations identified in the population were based on the degree of sequence conservation that is often poorly constructed. Second, our derivation of deleterious mutations does not include noncoding or structural variants, which can contribute considerably to the total load of mutations [53,54]. Third, our burden estimation assumes equal fitness effects for all mutations, which is unlikely, as mutations can have different fitness effects that can vary with environments [55]. Fourth, we considered the same sign of the effect when estimating the burden, which would be misestimated, as some deleterious mutations may be locally adaptive, or neutral [53,56]. Nonetheless, despite these caveats, our findings revealed a substantial genomic burden of deleterious mutations in sorghum.

We investigated the phenotypic effects of deleterious mutations (Table S1). We found negative correlations between mutation burden and phenotypic traits, suggesting a considerable cost of deleterious mutations on phenotypic traits [33] in a species that has been subjected to recent demographic expansion [50]. Consistently, we find a minor proportion of deleterious mutations with demonstrably large biological effects, which likely have an impact on phenotypes (Fig. 2). The fate of such large effect mutations on phenotypes is, however, unclear and has been actively debated as to whether such mutational effects primarily attributable to unconditional deleteriousness or can provide adaptive heritable variation [57]. Nonetheless, previous studies revealed that post-domestication mutations resulted in novel variations of genes in sorghum, and that neodiversity contributed to new adaptations in sorghum [57,58].

Across four traits, we find that putatively deleterious alleles explain roughly half of the genetic variance (46%-54%), but there is only a moderate improvement in total heritable variance explained for biomass (7.6%) and plant height (5.2%). Additionally, there is no advantage for SLA and starch (Fig 4). Such a difference in the contribution of deleterious alleles to traits was recently observed in maize where dominance contributed substantially to grain yield while phenology traits appeared to be largely additive [33]. Though the effects of mutations being deleterious or compensatory depends greatly upon the genetic background into which that mutation is incorporated [13], the trivial contributions of mutations to SLA and starch indicate that such mutations could be either nearly neutral or negatively synergistic. Our results therefore support the proposition that deleterious mutational effects vary with phenotypic traits and are often larger for fitness-related quantitative traits, while they are unclear for traits that are not directly linked to fitness [59]. Fitness-related quantitative traits, which are expected to have a more complex genetic architecture, could potentially carry a higher polygenic mutation burden that could considerably affect phenotypes [60].

Such propositions are also in line with the longstanding understanding that fitness-linked quantitative traits that show directional dominance generally exhibit inbreeding depression [41,61,62], which is strongly associated with the degree of deleterious mutations in the genome [29].

Finally, although our study did not account for sampling error while estimating an individual deleterious variant effect, which is generally greater for rare variants [38], our heritability estimates are consistent with the prediction abilities of phenotypic traits. Our work, therefore, adds to ongoing GWP efforts exploring the cumulative effects of deleterious mutations on phenotypic diversity [13,33]. However, since rare deleterious variants are less correlated with each other and their associations greatly suffer from low statistical power [59,63], employing either gene- and/or family-based approaches [38,40,63], or leveraging the phenotypic patterns [53] in which deleterious mutations have recognizable phenotypic consequences would aid future studies in determining how rare deleterious mutations within an individual shape its phenotype [53].

## CONCLUSIONS

We used phenotypic and genomic data from different racial groups of sorghum to show that sorghum accumulates a considerable number of deleterious mutations in the genome. Mutation burden differed substantially among racial groups where it negatively correlated with phenotypes. Genomic selection models encompassing deleterious mutations show variable phenotypic predictions across traits and, given the relatively high level of population structure in sorghum, disentangling deleterious effects at the single variant level would take a tremendous amount of effort and recombination. Deleterious variants could be prioritized through work with intermediate phenotypes or with more extensive evolutionary analysis among closely related species. Both of these avenues, if combined with high throughput genome editing, could be used to systematically start removing deleterious variants from elite biomass sorghum.

## MATERIALS AND METHODS

### Plant material, field experiments and phenotypic data

For this study, a diversity panel with 869 biomass sorghum lines was assembled [34,64]. Although phenotypic data for the entire panel was collected, only a subset of 229 lines for which WGS data were available was included in the study.

Field experiments were conducted in Illinois during 2016 in an augmented block design that consisted of 960 4-row plots with a row length of 3 m, 1.5 m alleys and 0.76 m row spacing. All plots were arranged in 40 rows and 24 columns. Target density of plant population was approximately 270,368 plants ha^−1^and experiments were planted in late May and harvested in early October. Plant height was measured from the ground to the uppermost leaf whorl 16 weeks after planting and averaged across the plot. Biomass data was collected at harvesting using a 4-row Kemper head attached to the John Deere 5830 tractor. A plot sampler equipment with near infrad-red sensor (model 130S, RCI engineering) was used to measure wet weight of total biomass (kg) and to quantify biomass moisture (%) and starch (%) contents of plants [65] in the 2 middle rows of each 4-row plot. Biomass yield in dry metric tons per hectare was calculated as: dry metric tons per ha = total plot wet weight (kg) × (1 − plot moisture) / (plot area in square meter /10,000) [64].

To estimate specific leaf area (SLA), the youngest fully expanded leaf from two randomly selected plants of the middle two rows of each plot were excised just above the ligule 60 to 70 days after planting. Damaged leaves were avoided. Excised leaves were then re-cut under water, and the cut surface kept immersed. In the laboratory, three 1.6 cm leaf discs were collected from the middle of each leaf whilst avoiding the mid-rib. Leaf discs were immediately transferred to an oven set at 60°C for two weeks. The dry mass of leaf discs was determined, and SLA was expressed as the ratio of fresh leaf area to dry leaf mass (cm^2^ g^−1^).

### Statistical analysis of phenotypic data

Phenotypic data analysis was conducted according to experimental design, which consisted of a series of incomplete blocks connected through common checks. The following model was used to get best linear unbiased prediction (BLUPs) for all genotypes included in the field trial:

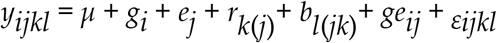

where *µ* is the overall mean, *g_i_* is the random effect of the *i^th^* genotype, *e_j_* is the random effect of the *j^th^* location, *r_k(j)_* is the random effect of the *k^th^* set nested within the *j^th^* location, *b_l(jk)_* is the random effect of the *l^th^* incomplete block nested within the *j^th^* location and the *k^th^* set, *g_eij_* represents the effect of genotype-by-environment interaction, and *ε_ijkl_* is the residual error for the *i^th^* genotype in the *l^th^* incomplete block within the *k^th^* set in the *j*^th^ location.

For the purpose of estimating the broad-sense heritability (*H^2^)* of each phenotype, we estimated variance component using the restricted maximum likelihood. All effects were assumed to be random effects. Broad-sense heritability on an entry-mean basis was calculated as H^2^ = σ^2^G / (σ^2^G + σ^2^GXE / number of location + σ^2^e / number of location × number of replication), where σ^2^G is the variance among accessions, σ^2^GXE is the accession-by-environment variance, and σ^2^e is the error variance. All analyses were conducted in R software (R Development Core Team, 2015) with package *lme4*.

### Genotyping

Genomic DNA (gDNA) was extracted using the CTAB method and quantified using picogreen (Molecular Probes, Eugene Oregon, USA) on a microplate reader of Synergy HT (BioTek, Vermont, USA). After preprocessing steps of the genomic DNA samples, ten libraries were prepared (24 samples in each library) and sequenced on HiSeq 4000 (PE_2×150) using sequencing kit version 1. Fastq files were demultiplexed with the bcl2fastq v2.17.1.14 conversion software of Illumina. We used Sentieon DNAseq [74] and a series of custom bash scripts to process the raw reads. Briefly, fastq files were aligned to the *Sorghum bicolor* reference genome version 3.1 (https://phytozome.jgi.doe.gov). PCR duplicates were removed, base quality was recalibrated based on a ‘known SNPs’ file, and recalibrated files were processed through the Haplotype Caller (HC). No realignment around indels was performed. The dataset therefore contains 239 samples, corresponding to 229 unique accessions, of which 7 had 1 or 2 replicates.

To create a list of “known SNPs” for the recalibration step, the HC pipeline was run without recalibration on the list of 239 BAM files. The output was filtered removing SNPs that had a number of heterozygote genotypes across all accessions greater than 10% and/or a number of heterozygote genotypes greater than two times the number of minor alleles (hereafter referred to as “homozygosity-based filter” [66]). In addition, “SNP clusters”, defined as three or more SNPs located within five base pairs (bp) were also filtered out. Clusters of SNPs are often generated by misalignment and were conservatively considered as spurious. The filtered list of SNPs was used as “known SNPs” to recalibrate the BAM files and to generate a final list of SNPs. To increase calibration accuracy, additional vcf files were also used as “known SNPs”. These vcf files were generated applying the same pipeline as described above to the publicly available fastq files of 42 *Sorghum bicolor* from [67] and an unpublished dataset of 302 *Sorghum bicolor* accessions kindly shared by Todd Mockler. The vcf file generated by the HC contained biallelic SNPs (*n*=22,359,733) and were further filtered to only retain SNPs with at least 4X coverage (*n*=21,865,512), and with a non-missing genotype in at least 40% of the samples (*n*=14,535,156). After removing SNPs clusters and applying homozygosity-based filters, the final dataset contained 5,512,653 SNPs, which were used for further analyses.

### Identifying putatively deleterious mutations

The substitution of amino acid effect on protein function was predicted with the SIFT algorithm [36]. A nonsynonymous mutation with a SIFT score <0.05 was defined as a putative deleterious mutation. In addition to SIFT, we also used genomic evolutionary rate profiling (GERP>2) [35] estimated from a multi-species whole-genome alignment of six species including *Zea mays*, *Oryza sativa*, *Setaria italica*, *Brachypodium distachyon*, *Hordeum vulgare*, and *Musa acuminate*. SIFT (<0.05) together with GERP (>2) annotations were combined to identify the deleterious mutations in constrained portions of the genome and defined as high GERP deleterious SNPs (HGERP_DEL-SNPs_). These mutations were used to estimate the mutation burden as the count of minor alleles present in each individual separately and corrected for the total number of non-missing sites within the individual [31]. Following Yang et al. [33], we defined the putative derived deleterious allele as a minor allele in the multi-species alignment.

We calculated linkage disequilibrium (LD) between SNPs to identify random variants (nondeleterious) to be used as a control to compare with deleterious mutations. A subset of 100k random SNP markers were selected, and all possible pairwise *r^2^* values were calculated using plink 1.9 [68]. Then, 1% of all the possible pairwise calculations were plotted showing the relationship of distance between markers and *r^2^*. To define local LD structure across each chromosome, we also calculated the mean LD score [69] per marker. LD scores were calculated with a window of 1Mb using the software GCTA [69,70]. Each LD score was divided by the total number of SNPs within each window (Fig. S6). To identify SNPs in high LD with deleterious variants, we first explored the effect of windows size and *r^2^* threshold on the number of SNPs selected (Fig. S7). Given the LD pattern observed, we used a window size of 250 kb and an *r^2^* threshold of 0.9, meaning that if any marker within 250 kb of a deleterious variants has an *r^2^* of 0.9 or higher, it would be excluded from further analysis. This yielded a list of ~1 million SNPs that were in LD with deleterious SNPs, which were excluded from all SNPs. An equal proportion of 100 sets of random variants with the similar allele frequency of deleterious variants were selected (Fig. S8).

### Estimating effect sizes of deleterious and common variants

Effect sizes were estimated using the RR-BLUP model implemented in the R-package rrBLUP version 4.2 [71]. We fit a model y = 1µ + Zu + e, where y is a vector of BLUPs of phenotype; 1µ is an intercept vector; Z is an *n* x *p* incidence matrix (either deleterious or random variants) containing the allelic states of the *p* marker loci ( *z* = {−1, 0, 1}), where −1 represents the minor allele; u is the *p* x 1 vector of marker effects; and e is a n × 1 vector of residuals. Under RR-BLUP, u ~ MVN (0, Iσ^2^_u_) where σ^2^ u is the variance of the common distribution of marker effects and was estimated using restricted maximum likelihood.

### Partitioning of genetic variance and genome-wide prediction

We compare the variance explained by deleterious variants to that of an equal proportion of randomly sampled variants from the distribution of non-deleterious variants. Following the method of Brenton et al. [34], we used a two-dimensional sampling approach to create 100 equal-sized datasets of randomly sampled variants matched for minor allele frequency. For each trait, we fit the model separately for each of variant set (either deleterious variant or non-deleterious variant) and estimated phenotypic variance explained.

For each variant set (deleterious variant vs non-deleterious set), we fit a standard GBLUP model including only additive effects by fitting a linear mixed model of the following form: *y* = Z*g* + e, where *y* is a vector of BLUPs of phenotype, the vector *g* is a random effect, the BLUP, which represents the GEBV for each individual, and Z is a design matrix indicating observations of genotype identities, and e is a vector of residuals. The genomic estimated breeding values (GEBV) were obtained by assuming g ~ MVN (0, Kσ^2^_g_), where σ^2^_g_ is the additive genetic variance, and K is the square genomic relationship matrix based on SNP data, implemented in TASSEL [72]. Predictive abilities for all traits were evaluated using a five-fold cross-validation approach repeated 100 times and were implemented in the R statistical software.

## ACKNOWLEDGEMENTS

The information, data, or work presented herein was funded in part by the Advanced Research Projects Agency-Energy (ARPA-E), U.S. Department of Energy, under Award Numbers DE-AR0000598 and DE-AR0000661. The views and opinions of authors expressed herein do not necessarily state or reflect those of the United States Government or any agency thereof. The support from the U.S. Department of Agriculture, Agricultural Research Service is greatly acknowledged.

## Supplementary Information

**Fig S1.**
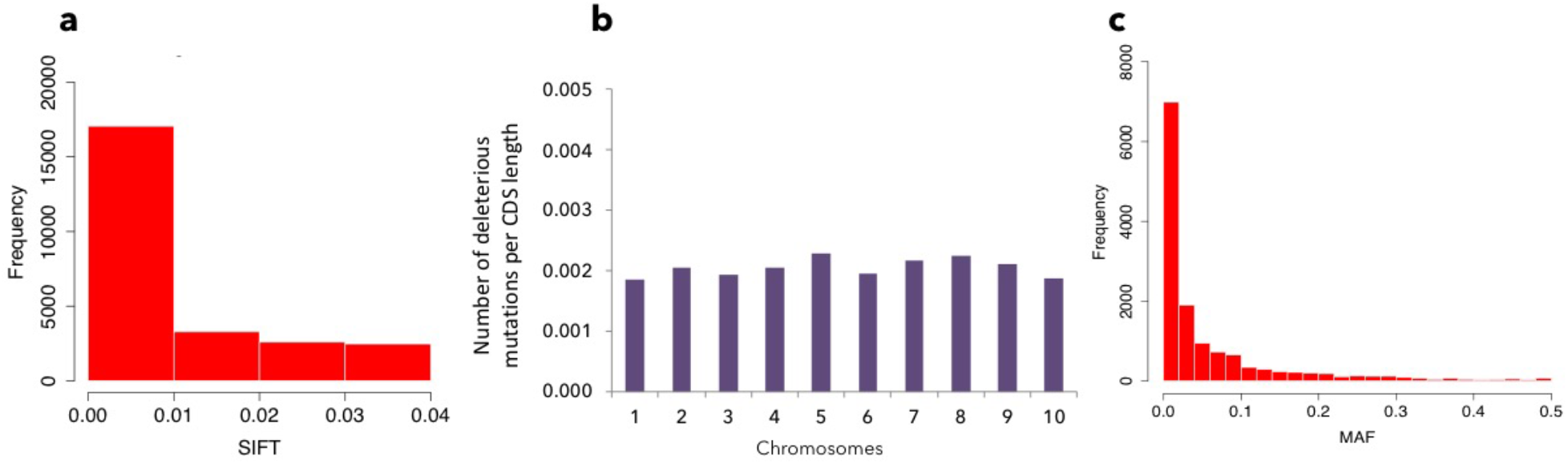
Frequency distributions of deleterious mutations. Sorting Intolerant From Tolerant (SIFT) distributions of (a) all deleterious mutations, (b) number of deleterious mutations estimated per coding regions in all chromosomes, and (c) allele frequency distribution of deleterious mutations.

**Fig S2.**
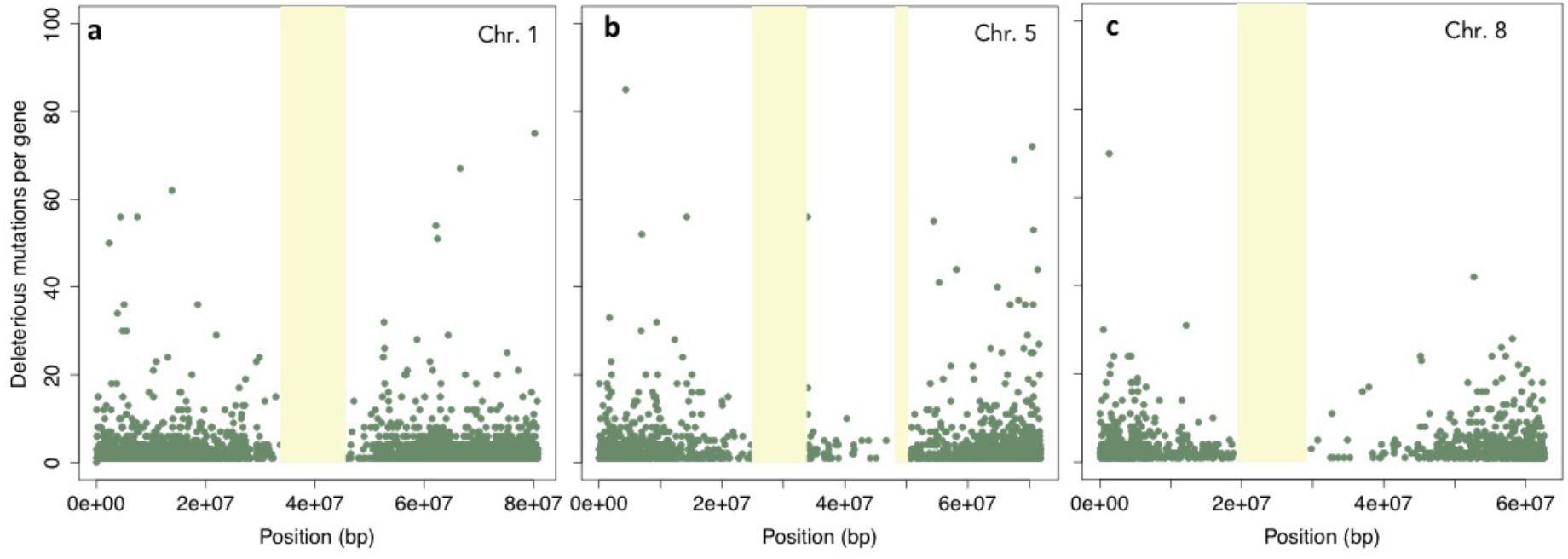
Gene deleterious mutations distribution in chromosomes 1 (a), 5 (b), and 8 (c). The yellow color vertical bar indicates a centromeric regions showing absence of genes or deleterious mutations.

**Fig S3.**
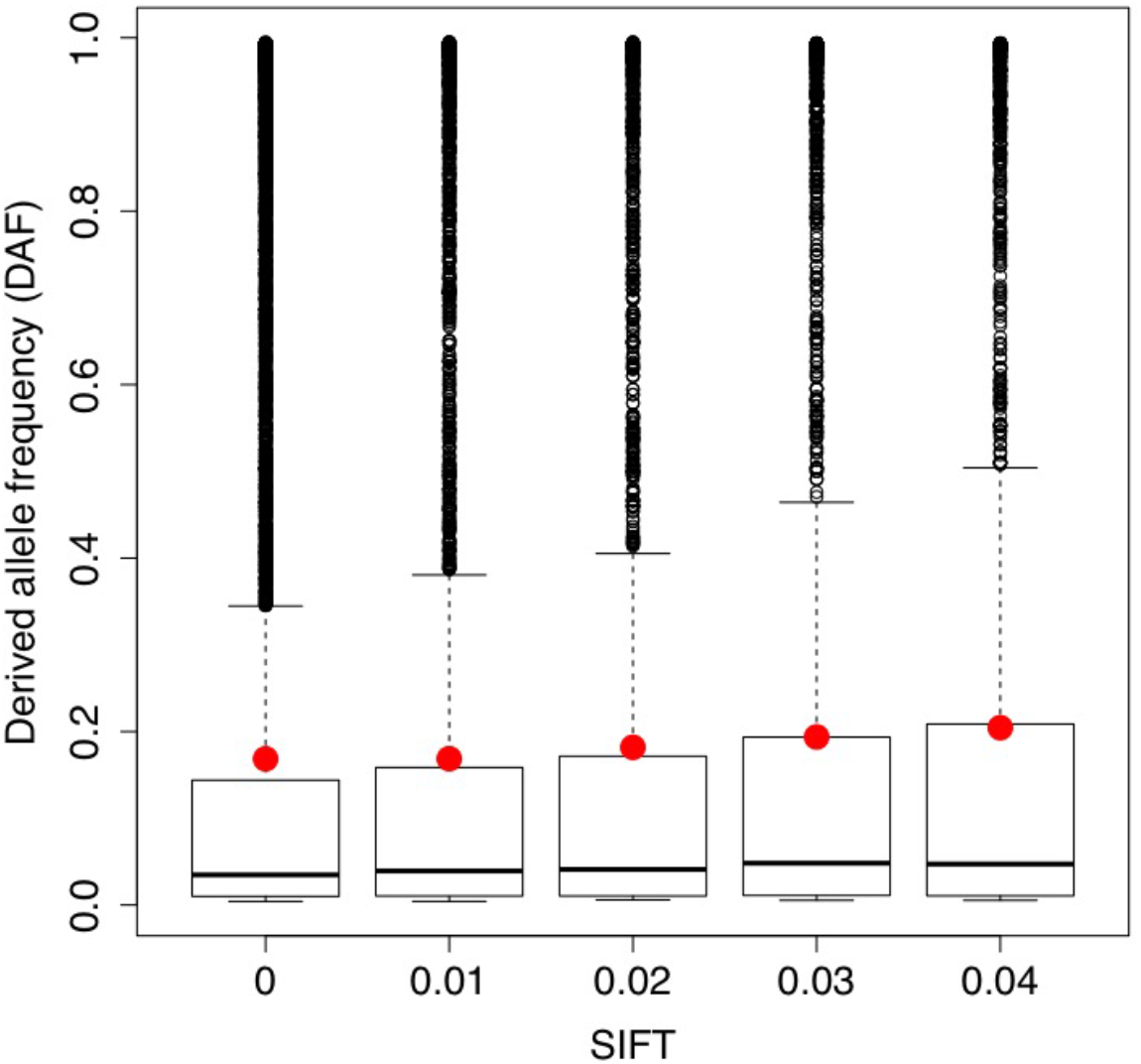
Derived allele frequency (DAF) association with Sorting Intolerant From Tolerant (SIFT) scores, Derived allele was defined as a minor allele from a multi-species sequence alignments. SIFT was estimated using [41].

**Fig S4.**
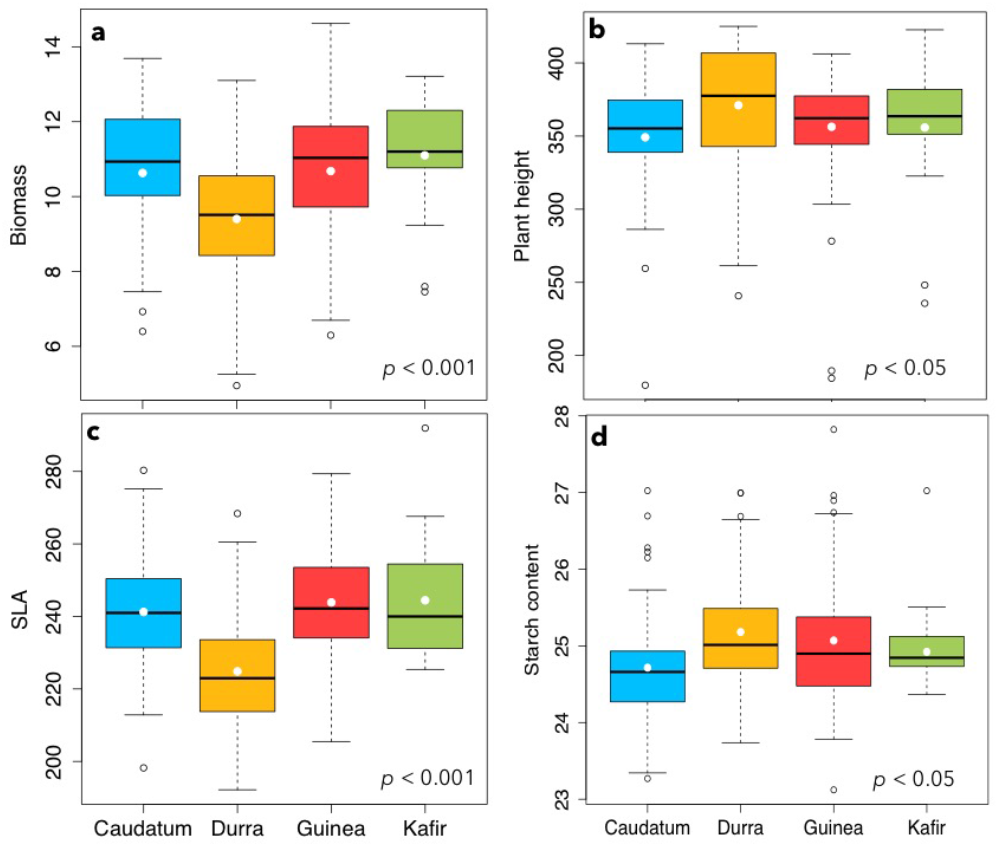
Boxplots of phenotypic data for biomass (a), plant height (b), specific leaf area (SLA, c), and tissue starch content (d) under different subpopulations of sorghum.

**Fig S5.**
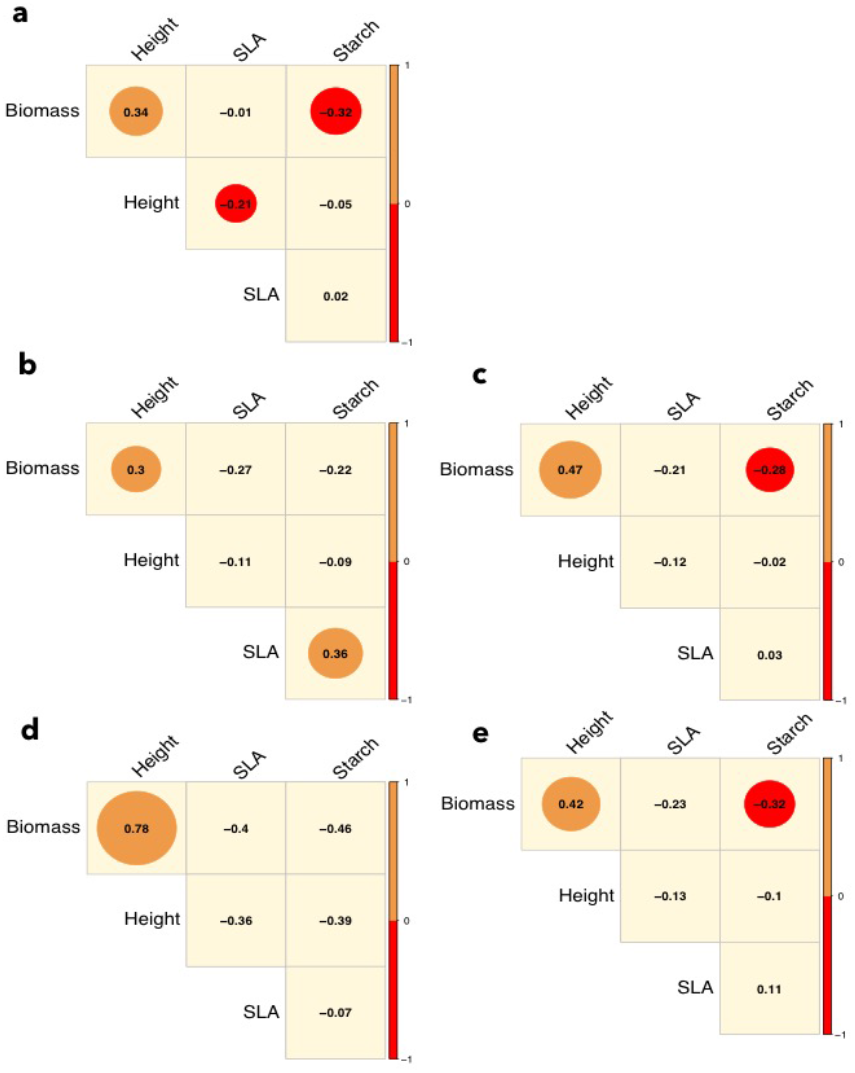
Correlations among traits either across all subpopulations (a) or within each subpopulation of durra (b), caudatum (c), kafir (d), and guinea (e) subpopulations. All circles around values indicates significance at P < 0.05. Orange circles indicate positive correlation while red circle represents a negative correlation.

**Fig S6.**
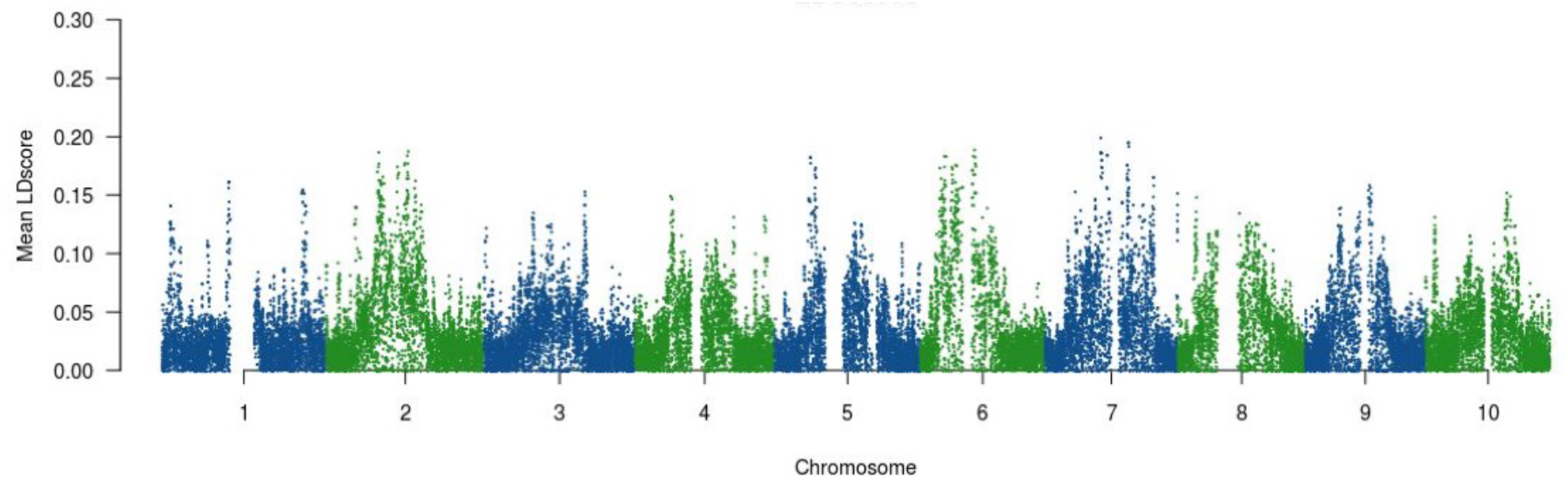
Mean linkage disequilibrium (LD) scores estimated for all chromosomes.

**Fig S7.**
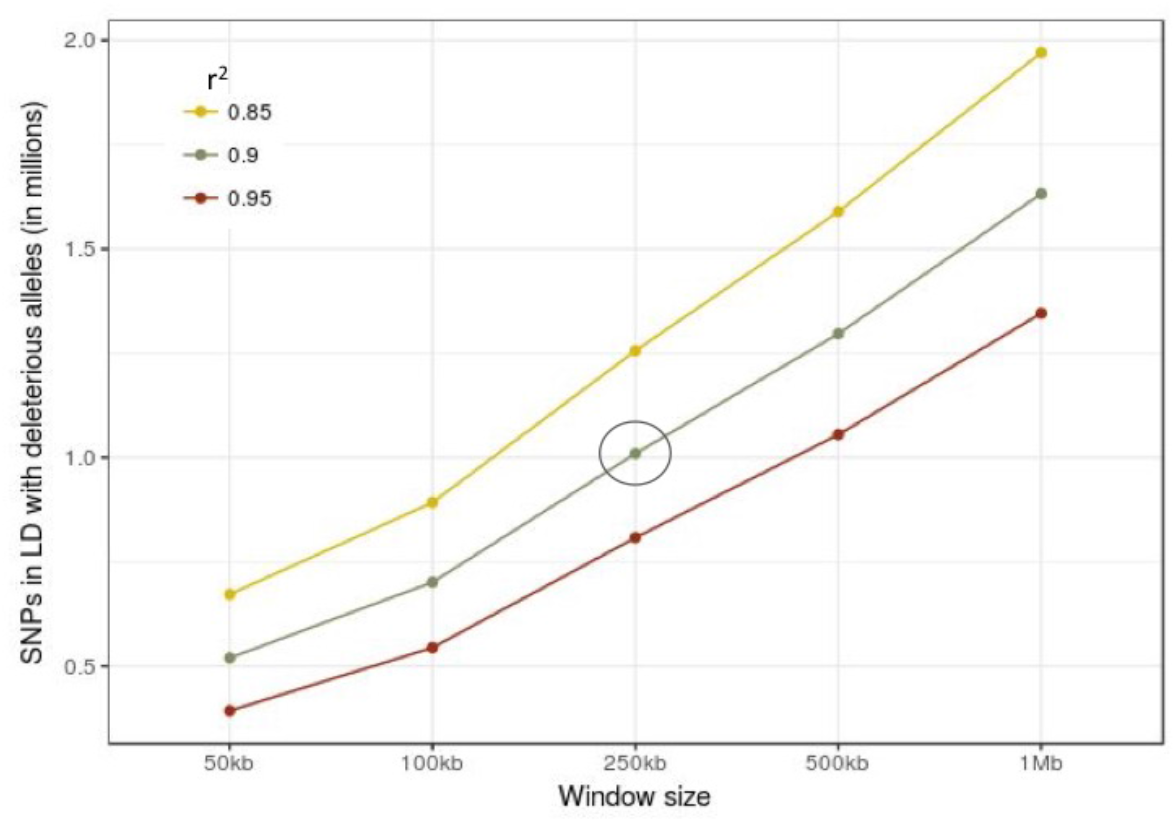
The number of single-nucleotide polymorphisms (SNPs) estimated under different parameters of window size and *r^2^*.

**Fig S8.**
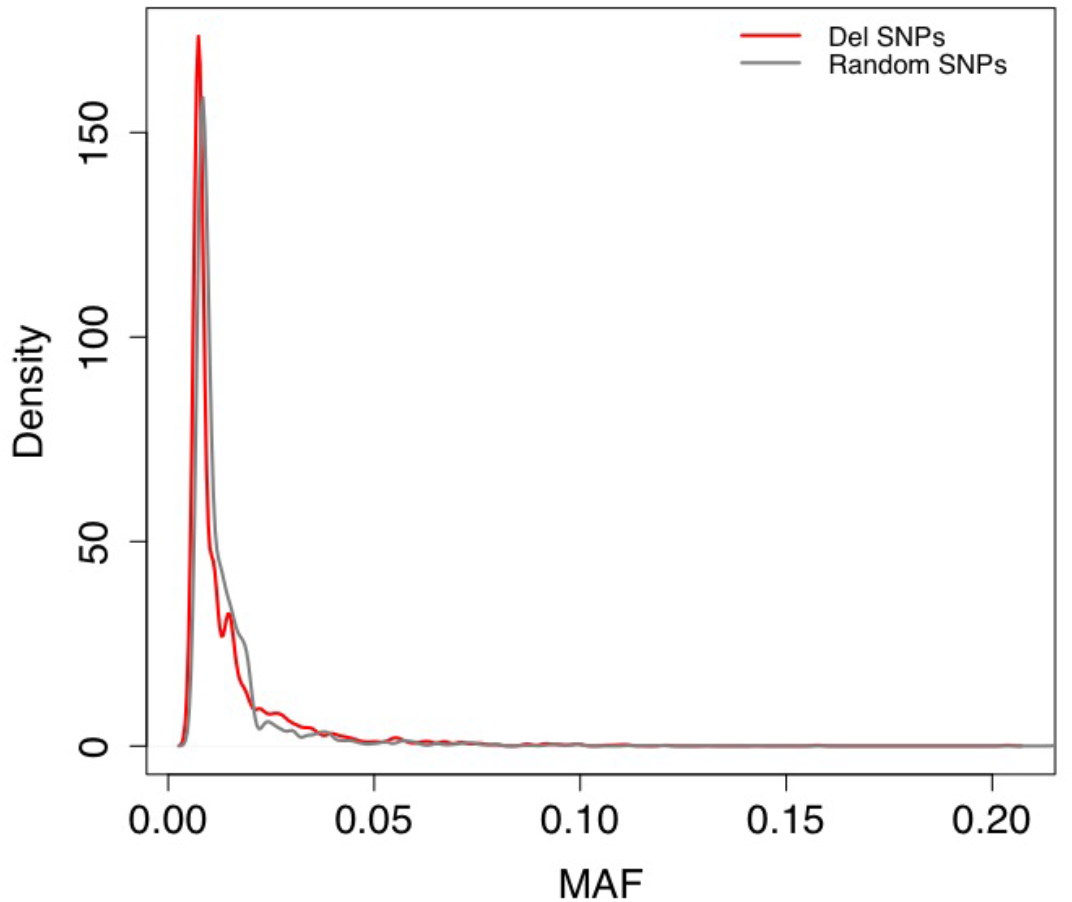
Allele frequency distribution comparison for deleterious mutations (red) and non-deleterious variants (grey).

**Table S1.** Slopes estimated between mutation burden and phenotypic traits.

| Trait | Estimate | $r^2$ | $P$ value |
| --- | --- | --- | --- |
| Biomass | -17.22 | 0.004 | 0.367 |
| Height | -71.27 | 0.014 | 0.081 |
| SLA | -18.55 | 0.005 | 0.279 |
| Starch | -11.60 | 0.010 | 0.153 |

## REFERENCES

1. Brandvain Y, Slotte T, Hazzouri KM, Wright SI, Coop G. Genomic Identification of Founding Haplotypes Reveals the History of the Selfing Species Capsella rubella. PLoS Genet. 2013;9. doi:10.1371/journal.pgen.1003754

2. Lynch M, Gabriel W. MUTATION LOAD AND THE SURVIVAL OF SMALL POPULATIONS. Evol Int J Org Evol. 1990;44: 1725–1737. doi:10.1111/j.1558-5646.1990.tb05244.x

3. Hartfield M, Glémin S. Hitchhiking of Deleterious Alleles and the Cost of Adaptation in Partially Selfing Species. Genetics. 2014;196: 281–293. doi:10.1534/genetics.113.158196

4. Ramu P, Esuma W, Kawuki R, Rabbi IY, Egesi C, Bredeson JV, et al. Cassava haplotype map highlights fixation of deleterious mutations during clonal propagation. Nat Genet. 2017;49: 959–963. doi:10.1038/ng.3845

5. Lu J, Tang T, Tang H, Huang J, Shi S, Wu C-I. The accumulation of deleterious mutations in rice genomes: a hypothesis on the cost of domestication. Trends Genet TIG. 2006;22: 126–131. doi:10.1016/j.tig.2006.01.004

6. Amorim CEG, Gao Z, Baker Z, Diesel JF, Simons YB, Haque IS, et al. The population genetics of human disease: The case of recessive, lethal mutations. PLOS Genet. 2017;13: e1006915. doi:10.1371/journal.pgen.1006915

7. Felsenstein J. The Evolutionary Advantage of Recombination. Genetics. 1974;78: 737–756.

8. Covert AW, Lenski RE, Wilke CO, Ofria C. Experiments on the role of deleterious mutations as stepping stones in adaptive evolution. Proc Natl Acad Sci. 2013;110: E3171–E3178. doi:10.1073/pnas.1313424110

9. Shaw FH, Geyer CJ, Shaw RG. A Comprehensive Model of Mutations Affecting Fitness and Inferences for Arabidopsis Thaliana. Evolution. 2002;56: 453–463. doi:10.1111/j.0014-3820.2002.tb01358.x

10. Doniger SW, Kim HS, Swain D, Corcuera D, Williams M, Yang S-P, et al. A Catalog of Neutral and Deleterious Polymorphism in Yeast. PLOS Genet. 2008;4: e1000183. doi:10.1371/journal.pgen.1000183

11. Yampolsky LY, Kondrashov FA, Kondrashov AS. Distribution of the strength of selection against amino acid replacements in human proteins. Hum Mol Genet. 2005;14: 3191–3201. doi:10.1093/hmg/ddi350

12. Kremling KAG, Chen S-Y, Su M-H, Lepak NK, Romay MC, Swarts KL, et al. Dysregulation of expression correlates with rare-allele burden and fitness loss in maize. Nature. 2018; doi:10.1038/nature25966

13. Moyers BT, Morrell PL, McKay JK. Genetic Costs of Domestication and Improvement. J Hered. 2018;109: 103–116. doi:10.1093/jhered/esx069

14. Wendorf F, Close AE, Schild R, Wasylikowa K, Housley RA, Harlan JR, et al. Saharan exploitation of plants 8,000 years BP. Nature. 1992;359: 721. doi:10.1038/359721a0

15. Dillon SL, Shapter FM, Henry RJ, Cordeiro G, Izquierdo L, Lee LS. Domestication to Crop Improvement: Genetic Resources for Sorghum and Saccharum (Andropogoneae). Ann Bot. 2007;100: 975–989. doi:10.1093/aob/mcm192

16. Evans J, McCormick RF, Morishige D, Olson SN, Weers B, Hilley J, et al. Extensive Variation in the Density and Distribution of DNA Polymorphism in Sorghum Genomes. PLOS ONE. 2013;8: e79192. doi:10.1371/journal.pone.0079192

17. Paterson AH, Bowers JE, Bruggmann R, Dubchak I, Grimwood J, Gundlach H, et al. The Sorghum bicolor genome and the diversification of grasses. Nature. 2009;457: 551. doi:10.1038/nature07723

18. Paterson AH, Bowers JE, Chapman BA. Ancient polyploidization predating divergence of the cereals, and its consequences for comparative genomics. Proc Natl Acad Sci U S A. 2004;101: 9903–9908. doi:10.1073/pnas.0307901101

19. Kumaravadivel N, Rangasamy SRS. Plant regeneration from sorghum anther cultures and field evaluation of progeny. Plant Cell Rep. 1994;13: 286–290. doi:10.1007/BF00233321

20. Arunkumar R, Ness RW, Wright SI, Barrett SCH. The Evolution of Selfing Is Accompanied by Reduced Efficacy of Selection and Purging of Deleterious Mutations. Genetics. 2015;199: 817–829. doi:10.1534/genetics.114.172809

21. Nakayama S-I, Shi S, Tateno M, Shimada M, Takahasi KR. Mutation Accumulation in a Selfing Population: Consequences of Different Mutation Rates between Selfers and Outcrossers. PLOS ONE. 2012;7: e33541. doi:10.1371/journal.pone.0033541

22. Pamilo P, Nei M, Li W-H. Accumulation of mutations in sexual and asexual populations. Genet Res. 1987;49: 135–146. doi:10.1017/S0016672300026938

23. Bustamante CD, Nielsen R, Sawyer SA, Olsen KM, Purugganan MD, Hartl DL. The cost of inbreeding in Arabidopsis. Nature. 2002;416: 531–534. doi:10.1038/416531a

24. Slotte T, Hazzouri KM, Ågren JA, Koenig D, Maumus F, Guo Y-L, et al. The *Capsella rubella* genome and the genomic consequences of rapid mating system evolution. Nat Genet. 2013;45: 831. doi:10.1038/ng.2669

25. Slotte T, Foxe JP, Hazzouri KM, Wright SI. Genome-wide evidence for efficient positive and purifying selection in Capsella grandiflora, a plant species with a large effective population size. Mol Biol Evol. 2010;27: 1813–1821. doi:10.1093/molbev/msq062

26. Gaut BS, Díez CM, Morrell PL. Genomics and the Contrasting Dynamics of Annual and Perennial Domestication. Trends Genet. 2015;31: 709–719. doi:10.1016/j.tig.2015.10.002

27. Kono TJY, Fu F, Mohammadi M, Hoffman PJ, Liu C, Stupar RM, et al. The Role of Deleterious Substitutions in Crop Genomes. Mol Biol Evol. 2016;33: 2307–2317. doi:10.1093/molbev/msw102

28. Günther T, Schmid KJ. Deleterious amino acid polymorphisms in Arabidopsis thaliana and rice. Theor Appl Genet. 2010;121: 157–168. doi:10.1007/s00122-010-1299-4

29. Mezmouk S, Ross-Ibarra J. The Pattern and Distribution of Deleterious Mutations in Maize. G3 Genes Genomes Genet. 2014;4: 163–171. doi:10.1534/g3.113.008870

30. Renaut S, Rieseberg LH. The Accumulation of Deleterious Mutations as a Consequence of Domestication and Improvement in Sunflowers and Other Compositae Crops. Mol Biol Evol. 2015;32: 2273–2283. doi:10.1093/molbev/msv106

31. Vitezica ZG, Varona L, Elsen J-M, Misztal I, Herring W, Legarra A. Genomic BLUP including additive and dominant variation in purebreds and F1 crossbreds, with an application in pigs. Genet Sel Evol. 2016;48: 6. doi:10.1186/s12711-016-0185-1

32. Vitezica ZG, Varona L, Legarra A. On the Additive and Dominant Variance and Covariance of Individuals Within the Genomic Selection Scope. Genetics. 2013;195: 1223–1230. doi:10.1534/genetics.113.155176

33. Yang J, Mezmouk S, Baumgarten A, Buckler ES, Guill KE, McMullen MD, et al. Incomplete dominance of deleterious alleles contributes substantially to trait variation and heterosis in maize. PLOS Genet. 2017;13: e1007019. doi:10.1371/journal.pgen.1007019

34. Brenton ZW, Cooper EA, Myers MT, Boyles RE, Shakoor N, Zielinski KJ, et al. A Genomic Resource for the Development, Improvement, and Exploitation of Sorghum for Bioenergy. Genetics. 2016;204: 21–33. doi:10.1534/genetics.115.183947

35. Habier D, Fernando RL, Kizilkaya K, Garrick DJ. Extension of the bayesian alphabet for genomic selection. BMC Bioinformatics. 2011;12: 186. doi:10.1186/1471-2105-12-186

36. Vaser R, Adusumalli S, Leng SN, Sikic M, Ng PC. SIFT missense predictions for genomes. Nat Protoc. 2016;11: 1. doi:10.1038/nprot.2015.123

37. Zöllner S, Pritchard JK. Overcoming the Winner’s Curse: Estimating Penetrance Parameters from Case-Control Data. Am J Hum Genet. 2007;80: 605–615.

38. Jun G, Manning A, Almeida M, Zawistowski M, Wood AR, Teslovich TM, et al. Evaluating the contribution of rare variants to type 2 diabetes and related traits using pedigrees. Proc Natl Acad Sci. 2018;115: 379–384. doi:10.1073/pnas.1705859115

39. Marouli E, Graff M, Medina-Gomez C, Lo KS, Wood AR, Kjaer TR, et al. Rare and low-frequency coding variants alter human adult height. Nature. 2017;542: 186–190. doi:10.1038/nature21039

40. Ji X, Kember RL, Brown CD, Bućan M. Increased burden of deleterious variants in essential genes in autism spectrum disorder. Proc Natl Acad Sci. 2016;113: 15054–15059. doi:10.1073/pnas.1613195113

41. Wright S. Evolution in Mendelian Populations. Genetics. 1931;16: 97–159.

42. Doggett H. Sorghum [Internet]. [London]: Longmans; 1970. Available: https://trove.nla.gov.au/version/45753139.

43. Lohmueller KE, Indap AR, Schmidt S, Boyko AR, Hernandez RD, Hubisz MJ, et al. Proportionally more deleterious genetic variation in European than in African populations. Nature. 2008;451: 994–997. doi:10.1038/nature06611

44. Fu W, Gittelman RM, Bamshad MJ, Akey JM. Characteristics of Neutral and Deleterious Protein-Coding Variation among Individuals and Populations. Am J Hum Genet. 2014;95: 421–436. doi:10.1016/j.ajhg.2014.09.006

45. Simons YB, Turchin MC, Pritchard JK, Sella G. The deleterious mutation load is insensitive to recent population history. Nat Genet. 2014;46: 220–224. doi:10.1038/ng.2896

46. Harlan JR, De Wet JMJ, Stemler ABL, ebrary I, Anthropological IC of, Ethnological Sciences (9th: 1973: Chicago I. Origins of African plant domestication [Internet]. The Hague: Mouton; Chicago: distributed by Aldine; 1976. Available: https://trove.nla.gov.au/version/13111119.

47. Stemler ABL, Harlan JR, Wet JMJ de. Evolutionary History of Cultivated Sorghums (Sorghum bicolor [Linn.] Moench) of Ethiopia. Bull Torrey Bot Club. 1975;102: 325–333. doi:10.2307/2484758

48. Ranwez V, Serra A, Pot D, Chantret N. Domestication reduces alternative splicing expression variations in sorghum. PLOS ONE. 2017;12: e0183454. doi:10.1371/journal.pone.0183454

49. Barro-Kondombo C, Sagnard F, Chantereau J, Deu M, Vom Brocke K, Durand P, et al. Genetic structure among sorghum landraces as revealed by morphological variation and microsatellite markers in three agroclimatic regions of Burkina Faso. TAG Theor Appl Genet Theor Angew Genet. 2010;120: 1511–1523. doi:10.1007/s00122-010-1272-2

50. Hamblin MT, Casa AM, Sun H, Murray SC, Paterson AH, Aquadro CF, et al. Challenges of Detecting Directional Selection After a Bottleneck: Lessons From Sorghum bicolor. Genetics. 2006;173: 953–964. doi:10.1534/genetics.105.054312

51. Szövényi P, Devos N, Weston DJ, Yang X, Hock Z, Shaw JA, et al. Efficient Purging of Deleterious Mutations in Plants with Haploid Selfing. Genome Biol Evol. 2014;6: 1238–1252. doi:10.1093/gbe/evu099

52. Charlesworth D, Wright SI. Breeding systems and genome evolution. Curr Opin Genet Dev. 2001;11: 685–690.

53. Bastarache L, Hughey JJ, Hebbring S, Marlo J, Zhao W, Ho WT, et al. Phenotype risk scores identify patients with unrecognized Mendelian disease patterns. Science. 2018;359: 1233–1239. doi:10.1126/science.aal4043

54. Huang Y-F, Gulko B, Siepel A. Fast, scalable prediction of deleterious noncoding variants from functional and population genomic data. Nat Genet. 2017; advance online publication. doi:10.1038/ng.3810

55. Henn BM, Botigué LR, Peischl S, Dupanloup I, Lipatov M, Maples BK, et al. Distance from sub-Saharan Africa predicts mutational load in diverse human genomes. Proc Natl Acad Sci. 2016;113: E440–E449. doi:10.1073/pnas.1510805112

56. Vikram P, Swamy BPM, Dixit S, Singh R, Singh BP, Miro B, et al. Drought susceptibility of modern rice varieties: an effect of linkage of drought tolerance with undesirable traits. Sci Rep. 2015;5: 14799. doi:10.1038/srep14799

57. Glémin S, Bataillon T. A comparative view of the evolution of grasses under domestication. New Phytol. 2009;183: 273–290. doi:10.1111/j.1469-8137.2009.02884.x

58. Figueiredo LF de A, Calatayud C, Dupuits C, Billot C, Rami J-F, Brunel D, et al. Phylogeographic Evidence of Crop Neodiversity in Sorghum. Genetics. 2008;179: 997–1008. doi:10.1534/genetics.108.087312

59. Park J-H, Gail MH, Weinberg CR, Carroll RJ, Chung CC, Wang Z, et al. Distribution of allele frequencies and effect sizes and their interrelationships for common genetic susceptibility variants. Proc Natl Acad Sci. 2011;108: 18026–18031. doi:10.1073/pnas.1114759108

60. Purcell SM, Moran JL, Fromer M, Ruderfer D, Solovieff N, Roussos P, et al. A polygenic burden of rare disruptive mutations in schizophrenia. Nature. 2014;506: 185–190. doi:10.1038/nature12975

61. Charlesworth B, Charlesworth D. The genetic basis of inbreeding depression. Genet Res. 1999;74: 329–340.

62. Kelly JK. An experimental method for evaluating the contribution of deleterious mutations to quantitative trait variation. Genet Res. 1999;73: 263–273.

63. Auer PL, Lettre G. Rare variant association studies: considerations, challenges and opportunities. Genome Med. 2015;7. doi:10.1186/s13073-015-0138-2

64. Fernandes SB, Dias KOG, Ferreira DF, Brown PJ. Efficiency of multi-trait, indirect, and trait-assisted genomic selection for improvement of biomass sorghum. Theor Appl Genet. 2018;131: 747–755. doi:10.1007/s00122-017-3033-y

65. Li J, Danao M-GC, Chen S-F, Li S, Singh V, Brown PJ. Prediction of Starch Content and Ethanol Yields of Sorghum Grain Using near Infrared Spectroscopy. J Infrared Spectrosc. 2015;23: 85–92.

66. Chia J-M, Song C, Bradbury PJ, Costich D, de Leon N, Doebley J, et al. Maize HapMap2 identifies extant variation from a genome in flux. Nat Genet. 2012;44: 803–807. doi:10.1038/ng.2313

67. Mace ES, Tai S, Gilding EK, Li Y, Prentis PJ, Bian L, et al. Whole-genome sequencing reveals untapped genetic potential in Africa’s indigenous cereal crop sorghum. Nat Commun. 2013;4: 2320. doi:10.1038/ncomms3320

68. Chang CC, Chow CC, Tellier LC, Vattikuti S, Purcell SM, Lee JJ. Second-generation PLINK: rising to the challenge of larger and richer datasets. GigaScience. 2015;4: 7. doi:10.1186/s13742-015-0047-8

69. Bulik-Sullivan BK, Loh P-R, Finucane HK, Ripke S, Yang J, Schizophrenia Working Group of the Psychiatric Genomics Consortium, et al. LD Score regression distinguishes confounding from polygenicity in genome-wide association studies. Nat Genet. 2015;47: 291–295. doi:10.1038/ng.3211

70. Yang J, Lee SH, Goddard ME, Visscher PM. GCTA: a tool for genome-wide complex trait analysis. Am J Hum Genet. 2011;88: 76–82. doi:10.1016/j.ajhg.2010.11.011

71. Endelman JB. Ridge regression and other kernels for genomic selection with R package rrBLUP. The Plant Gen. 2011; 4:250–255. doi:10.3835/plantgenome2011.08.0024

72. Bradbury PJ, Zhang Z, Kroon DE, Casstevens TM, Ramdoss Y, Buckler ES. 2007. TASSEL: Software for association mapping of complex traits in diverse samples. Bioinformatics 23:2633–2635. doi.org/10.1093/bioinformatics/btm308

